# KnowMore: an automated knowledge discovery tool for the FAIR SPARC datasets

**DOI:** 10.1101/2021.08.08.455581

**Authors:** Ryan Quey, Matthew A. Schiefer, Anmol Kiran, Bhavesh Patel

## Abstract

This manuscript provides the methods and outcomes of KnowMore, the Grand Prize winning automated knowledge discovery tool developed by our team during the 2021 NIH SPARC FAIR Data Codeathon. The National Institutes of Health Stimulating Peripheral Activity to Relieve Conditions (NIH SPARC) program generates rich datasets from neuromodulation researches, curated according to the Findable, Accessible, Interoperable, and Reusable (FAIR) SPARC data standards. These datasets are publicly available through the SPARC Data Portal at sparc.science. Currently, the process of simultaneously comparing and analyzing multiple SPARC datasets is tedious because it requires investigating each dataset of interest individually and downloading all of them to conduct cross-analyses. It is crucial to enhance this process to enable rapid discoveries across SPARC datasets. To fill this need, we created KnowMore, a tool integrated into the SPARC Portal that only requires the user to select their datasets of interest to launch an automated discovery process. KnowMore uses several SPARC resources (Pennsieve, o^2^S^2^PARC, SciCrunch, protocols.io, Biolucida), data science methods, and machine learning algorithms in the back end to generate various visualizations in the front end intended to help the user identify potential similarities, differences, and relations across the datasets. These visualizations can lead to a new discovery, new hypothesis, or simply guide the user to the next logical step in their discovery process. The outcome of this project is a SPARC portal-ready code architecture that helps researchers to use SPARC datasets more efficiently and fully leverages their FAIR characteristics. The tool has been built and documented such that more data analysis methods and visualization items could be easily added. The potential for automated discoveries from SPARC datasets is huge given the unique SPARC data ecosystem promoting FAIR data practices, and KnowMore has only demonstrated a small highlight of what could be achieved to speed up discoveries from SPARC datasets.

## Introduction

The National Institutes of Health’s (NIH’s) Stimulating Peripheral Activity to Relieve Conditions (SPARC) program seeks to accelerate the development of therapeutic devices that modulate electrical activity in nerves to improve organ function^1^. A major focus of the SPARC program is to generate rich datasets that provide resources for understanding nerve-organ interactions and guiding the development of neuromodulation therapies. These datasets are publicly available through an open data platform, the SPARC Data Portal^2^. As of July 2021, 115 datasets are available spanning multiple scales (cellular, tissue, organ level), organs (stomach, large intestine, small intestine, heart, bladder, urinary tract, lung, pancreas, spleen), species (pig, human, rat, mouse, dog), and data types (scaffold data, histology, immunohistochemistry, electrical impedance tomography, 3D microscopy, morphometric analyses, computer simulations of single axons or populations of axons, electrophysiological responses to electrical stimulation, etc.).

To ensure SPARC datasets are findable, accessible, interoperable, and reusable (FAIR), they are curated according to the SPARC Data Structure (SDS), the data standards designed by the SPARC Data Curation Team to capture the large variety of data generated by SPARC investigators^3,4^. Accordingly, many resources are made available to SPARC researchers for making their data FAIR, such as the cloud data platform Pennsieve, the curated vocabulary selector and annotation platform SciCrunch, the open source computational modeling platform Open Online Simulations for Stimulating Peripheral Activity to Relieve Condition (o^2^S^2^PARC), the online microscopy image viewer Biolucida, and the data curation Software to Organize Data Automatically (SODA)^5,6^. As a result, the SPARC program provides a wealth of open and well-curated datasets that are accessible via the SPARC Data Portal. The portal provides several means of accessing data. A standard portal search feature is available. Alternatively, the user can find datasets by browsing through data categorized by organ system. The user can also use an interactive map to click on organs or nerves in animal models of interest and the portal provides links to associated datasets. These pathways make it easy to find datasets. Clicking a link to a dataset provides the user with details about the study and options to download all or subsets of the data files.

While it is very easy to look at the details of any single SPARC dataset on the portal, there is currently no easy way to rapidly compare multiple datasets. Typically, a researcher wanting to find relations across datasets would have to do so manually by going through each dataset individually, i.e., read the description of each dataset, go through each protocol, browse files that are accessible from the browser, etc. Datasets that warrant further investigation must be downloaded for offline analyses and payment may be required for access to large datasets, according to Amazon Web Services (AWS) pricing. Depending on the formats of the data, this may require programming skills beyond that of many users. After spending time collating data in a form that allows comparison across the different datasets, the user may find that, in fact, the datasets did not contain the information they needed. This process of analyzing multiple datasets together is tedious, which ideally should not be the case since the SDS is designed to facilitate such analysis. Therefore, this process needs to be urgently improved to 1) enable rapid discoveries across SPARC datasets and 2) encourage more researchers to use the SPARC Data Portal.

To address this shortcoming, we developed KnowMore during the 2021 SPARC FAIR Codeathon^7^ (July 12^th^, 2021 – July 26^th^, 2021). KnowMore is an automated knowledge discovery tool integrated within the SPARC Portal. With minimal clicks, the user selects datasets of interest and KnowMore allows the user to visualize potential relations, similarities, differences, and correlations between the studies and associated datasets. This process, illustrated in Figure 1, is achieved by leveraging our knowledge of the SPARC data structure and metadata that allows us to perform text mining, generate a summary table, and plot data that is common across all selected datasets. The results are presented as several visualization items that provide the user with a quick means of identifying potential relations across the datasets. This manuscript describes the structure of KnowMore and provides an example of knowledge provided by the tool when applied to a set of three sample datasets that constituted our use case for demonstrating KnowMore during the 2021 SPARC FAIR Codeathon.

**Figure 1.**
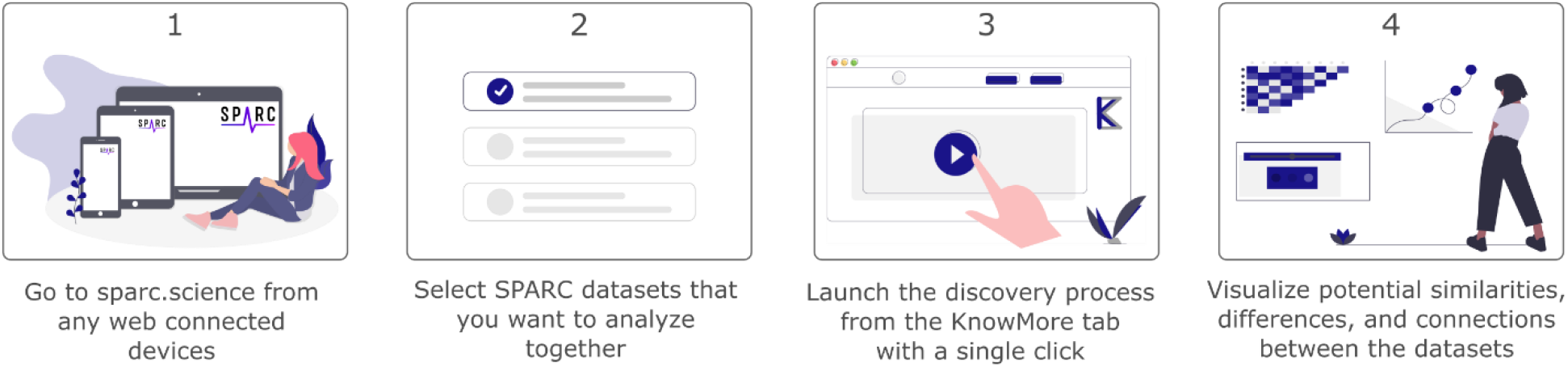
**Illustration of the simple user side workflow of KnowMore. Note that the tool is not currently integrated in the official SPARC Portal, but accessible through our fork of the sparc-app. It could be included into the official SPARC Portal after future consultation with SPARC**.

## Methods

### Software architecture

The overall workflow of KnowMore is shown in Figure 2. Our architecture consists of three main blocks that are independent:

**Figure 2.**
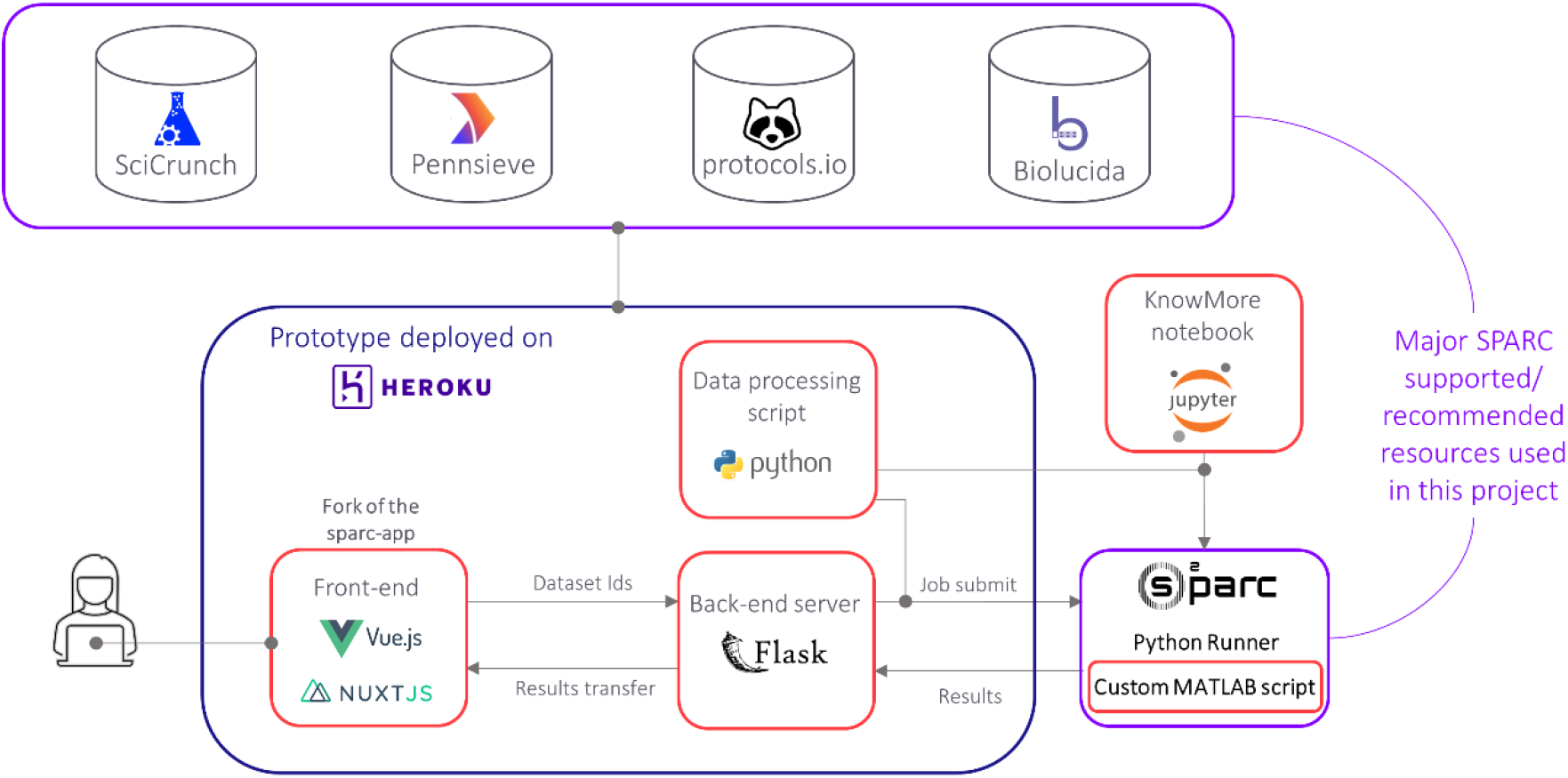
**Illustration of the overall technical workflow of KnowMore. The red rectangles highlight the major code blocks of KnowMore that were developed during the 2021 SPARC FAIR Codeathon**.

1. The front end of our app is based on a fork of the sparc-app (i.e. the front end of sparc.science) where we have integrated additional user interface elements and front end logic for KnowMore^8^.
2. The back end consists of a Flask (a micro web framework written in Python programming language) application that listens to front end requests and launches the data processing jobs.
3. The data processing and result generation are done through a MATLAB code (for ‘MAT’ data files) and a Python code (all other data types) that both run on the o^2^S^2^PARC platform, the SPARC supported cloud computing platform^9^.

In our front end of the sparc-app, we have integrated an “Add to KnowMore” button that is visible in the search result for each dataset and also available on the dataset page. By clicking on this button, the user can add their desired datasets for the analysis. Once all the datasets have been added, the user can go to the “KnowMore” tab we have included. On that page, the user can see a list of the selected datasets as well as a “Discover” button. A click on that button initiates the discovery process, where the Pennsieve IDs (i.e., the unique ID attributed to each dataset on sparc.science) of the selected datasets are sent to the Flask server, which then sends the IDs and our data processing Python script to o^2^S^2^PARC, using the o^2^S^2^PARC application programming interface (API)^10^. Once the script is fully executed, the results are sent back to the Flask server, which then transfers them to the front end where visualization items are generated. More details about the visualization items are provided in the next section.

The software architecture shown in Figure 2 was motivated by our aim of making KnowMore ready to on-board the SPARC Data Portal:

- Integrating the front end of KnowMore will only require merging our fork of the sparc-app with the main branch sparc-app.
- The back end of the sparc-app, the sparc-api, is built with Flask so the KnowMore back end is readily integrable^11^.
- The data processing jobs are designed to run on o^2^S^2^PARC and do not require any type of integration as our back end ensures communication with o^2^S^2^PARC.

Moreover, each of the three main elements of KnowMore is fully independent. While the front end will not be of much use on its own, having the back end fully interoperable is very valuable as our Flask application can be connected to any front end if needed (another analysis tool, website, software, etc.). The data processing and results generation jobs are also independent such that they can be used directly to get the visualization items. We have demonstrated that by developing a Jupyter Notebook that communicates directly with o^2^S^2^PARC to run the knowledge discovery jobs based on user-specified dataset IDs. Note that the data for the Knowledge Graph is obtained from Pennsieve/Scicrunch on the front end for efficiency but the same results can be generated in the back end as well. Thorough details for using the source code are available on the GitHub repository for this project^12^.

### Data processing and outputs

The output of KnowMore consists of multiple interactive visualization items displayed to the user so that they can progressively gain knowledge on the potential similarities, differences, and relations across the datasets. This output is intended to provide foundational information to the user so that they can rapidly make novel discoveries from SPARC datasets, generate new hypotheses, or simply decide on their next step (assess each dataset individually on the portal, download and analyze the datasets further, remove/add datasets to their analysis pool, etc.). A list of the visualization items is provided in Table 1, along with the potential knowledge that could be gained from each of them.

**Table 1.** Table listing the visualization items automatically generated by KnowMore. The source of the raw data for generating the visualization items are also listed.

| Visualization item | Knowledge gained across the datasets | Raw data used for generating the visualization and how it was obtained |
| --- | --- | --- |
| Knowledge Graph | High-level connections (authors, institutions, funding organisms, etc.) | Dataset metadata obtained with the Pennsieve API and the SciCrunch Elasticsearch API |
| Summary Table | Similarities/differences in the study design | Dataset metadata from the metadata.json file obtained with the Pennsieve API |
| Common Keywords | Common themes | Dataset metadata from the metadata.json file and all dataset text files obtained with the Pennsieve API. Protocol text obtained with the protocols.io API |
| Abstract | Common study design and findings | Dataset metadata from the metadata.json file and all dataset text files obtained with the Pennsieve API. Protocol text obtained with the protocols.io API |
| Data Plots | Comparison between measured numerical data (if any) | MAT files in the derivative folder of the datasets obtained with the Pennsieve API |

The process of getting these outputs starts by getting the IDs of the datasets selected by the user, which are obtained using the Pennsieve API^13^ in the front end. From there, we leverage several SPARC-supported and recommended resources in our data processing Python Script to collect the raw data required to generate the above-mentioned outputs. These resources include the Pennsieve API^13^, the Scicrunch Elasticsearch API^14^, the protocols.io API^15^, and the Biolucida API^16^. We refer to the paper on the SPARC Data Resource Center (DRC) for more details about these resources and their role in the SPARC data ecosystem^6^. Details about each of the visualization items are provided below. Each of these items can be easily saved from the front-end interface.

#### Knowledge graph

Using the Pennsieve ID of each dataset, the following items are queried from SciCrunch Elasticsearch API^14^: Person (authors of the dataset), Affiliation (affiliation of the authors), and Award (funding source for the dataset). The visualization library Vega is used in the front end to display this information in an interactive knowledge graph, which instantly highlights high-level relations amongst the datasets.

#### Summary table

A summary table is built with information collected from the metadata.json file of each dataset, which is a standard file generated for each SPARC dataset when published, and the subjects and samples metadata files, which are standard metadata files prescribed for SPARC datasets by the SDS. The files are retrieved from the Pennsieve API within our Python code. The following items are parsed from the metadata.json file for each dataset: title of the dataset, subtitle of the dataset, publication date. The following items are parsed from the subjects’ metadata file for each dataset: number of subjects, species, age, sex. The following items are parsed from the samples metadata file: number of samples, specimen, type, specimen anatomical locations. The visualization library Plotly is used in the front end to display the results in an interactive table, which shows this information side-by-side for each dataset, thus enabling quick comparison in the study design of each dataset.

#### Keywords

Text is obtained for each dataset from the description included in metadata.json file and the text from all the text files in the dataset using the Pennsieve API, and the text from the protocol on protocols.io associated with the dataset using the protocols.io API. The link to the protocol.io protocol is extracted from the metadata.json file of the dataset. All text is combined to create a paragraph for each dataset. The Natural Language Processing (NLP) Python library NLTK^17^ is then used to clean the text (e.g., remove stopwords). Biological keywords are identified using the spaCy python module and ScispaCy models^18,19^. The frequency of biological words is counted for each dataset. The final frequency of the keywords is assigned based on lowest occurrence among the datasets and the twenty most frequent words are selected and displayed as a word cloud using the visualization library Vega. The minimum frequency of a keyword across the dataset is displayed when the cursor hovers over the word. These keywords conveniently allow the user to identify common themes across the datasets.

#### Correlation matrix and abstract

The correlation matrix demonstrates the putative relatedness between datasets^20,21^. To generate the correlation matrix for the given datasets, pairwise similarity between datasets is calculated using the following Jaccard index equation^22^:

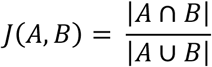

where A and B are sets of biological keywords present in two datasets. The biological keywords are identified as explained earlier in the Keywords section.

Paragraphs generated from datasets for the keywords identification are merged and divided into sentences. Each sentence is further divided into words and stopwords were removed. The frequency of each remaining word in a sentence is counted and converted into vectors where keywords represent the direction and frequencies represent the magnitude. The distance of two sentences is calculated using equation 1 − cos(*θ*) where cosine similarity is expressed as follows:

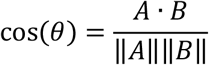

where A and B are words frequency in vectors of two sentences. Based on the pairwise distance of sentences a pagerank is assigned to each sentence using Python networkX module and sentences are ordered based on pagerank in decreasing order^23^. The top 10 highest-ranked sentences are selected to generate a common abstract for the datasets. This abstract is intended to provide a quick idea of any common study design and/or findings.

#### Data plots

If data files in .mat format are found under the “derivative” folder, the data processing Python script extracts and saves them then provides them to our MATLAB script that is compiled and deployed on o^2^S^2^PARC. The script collates the data into a data table. The script next determines which columns in the data table can be used for plotting purposes. Columns containing categorical data are limited to the x-axis. Columns containing numerical data can be plotted on the x-axis or y-axis. Columns containing any other type of data are excluded. Plots are then generated for every variable that can be displayed on a y-axis against every variable that can be plotted on an x-axis. In addition to the plots, the MATLAB script outputs an Excel file that lists each of the plots created and the variables included in each plot. The Excel file also includes data for each plot. Additionally, the script creates a json file that includes all data for each plot. These plots quickly highlight to the user relations between similar quantities measured across datasets.

#### Image clustering

An additional visualization item we aimed to provide to the user but could not complete during the Codeathon due to time constraints was a clustering of images across datasets, which may be particularly useful for histological data. All image data from SPARC datasets are stored on Biolucida. We currently have a function in place to retrieve image data from Biolucida given a Pennsieve dataset ID using the Biolucida API^16^. In the future, image clustering and visualization components will be added in the Python script and front end, respectively, to provide an additional element to the user for comparing datasets.

### Use case

#### Setup

KnowMore was developed and tested using three datasets available at sparc.science (Table 2). These datasets were selected because they have a common theme – quantified vagus nerve morphology – and span three species: rat, pig, and human. In principle, KnowMore is not specifically designed around these datasets and is coded to work with any user-selected datasets. However, for demonstration purposes, the data plots are currently limited to only appear when working with all or a subset of the three datasets listed in Table 2. Reasons for this are addressed in the Challenges section below and recommendations are put forth to expand the usability of this feature and increase the interoperability of SPARC datasets.

**Table 2.** List of datasets used for our use case.

| **Pennsieve ID** | **Title** |
| --- | --- |
| 60 | Quantified Morphology of the Rat Vagus Nerve<sup>25</sup> |
| 64 | Quantified Morphology of the Pig Vagus Nerve<sup>26</sup> |
| 65 | Quantified Morphology of the Human Vagus Nerve with Anti-Claudin-1<sup>27</sup> |

Initiating a KnowMore analysis requires five steps:

1. Use the search feature or browse for possible datasets of interest at sparc.science.
2. As datasets are identified that the user wants to compare, click on the “Add to KnowMore” button, visible in the header of the datasets or the search results. This will add the datasets to the KnowMore analysis.
3. Go to the KnowMore tab at the top of the webpage and check that all of the desired datasets are listed.
4. Decide which output to display. All possible output is displayed by default.
5. Click on the “Discover” button to initiate the automated analysis.

The number of datasets selected will affect the duration of time required to run the full discovery analysis. The use case with these three datasets takes about 4 min to generate all the visualization items.

### Outputs

#### Knowledge Graph

The Knowledge Graph provides an interactive tool to visualize metadata across the three datasets (Figure 3). This provides the ability to quickly determine, for example, that all three datasets had four investigators in common (Cariello, Grill, Goldhagen, and Pelot) affiliated with the Department of Biomedical Engineering at Duke and that the human dataset had additional investigators (Ezzell and Clissold) affiliated with the Department of Cell Biology and Physiology at the University of North Carolina.

**Figure 3.**
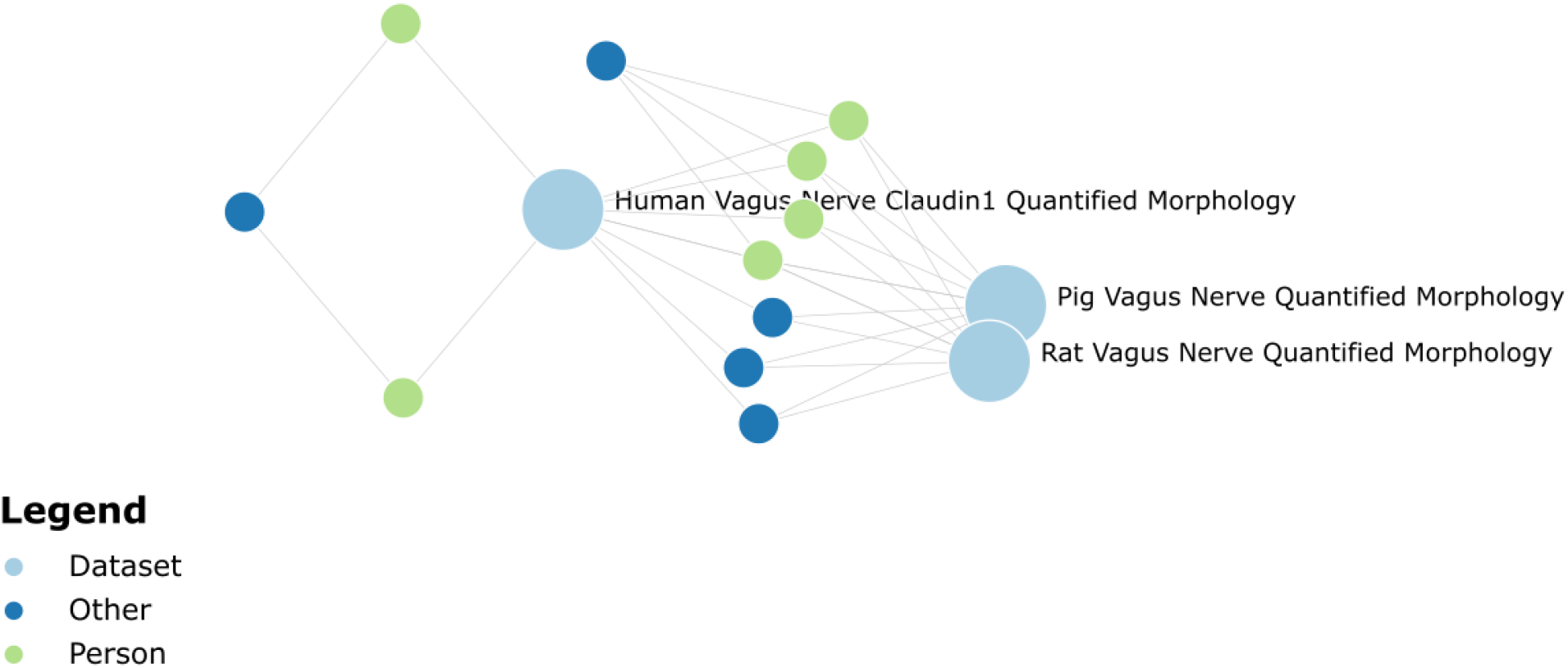
Knowledge Graph output for the three datasets in our use case.

#### Summary Table

The Summary Table provides the user with key pieces of information from each study in tabular format (Table 3). From this table, the user can easily determine that datasets have several common metrics. However, perineurial thickness is not quantified in dataset 64.

**Table 3.**
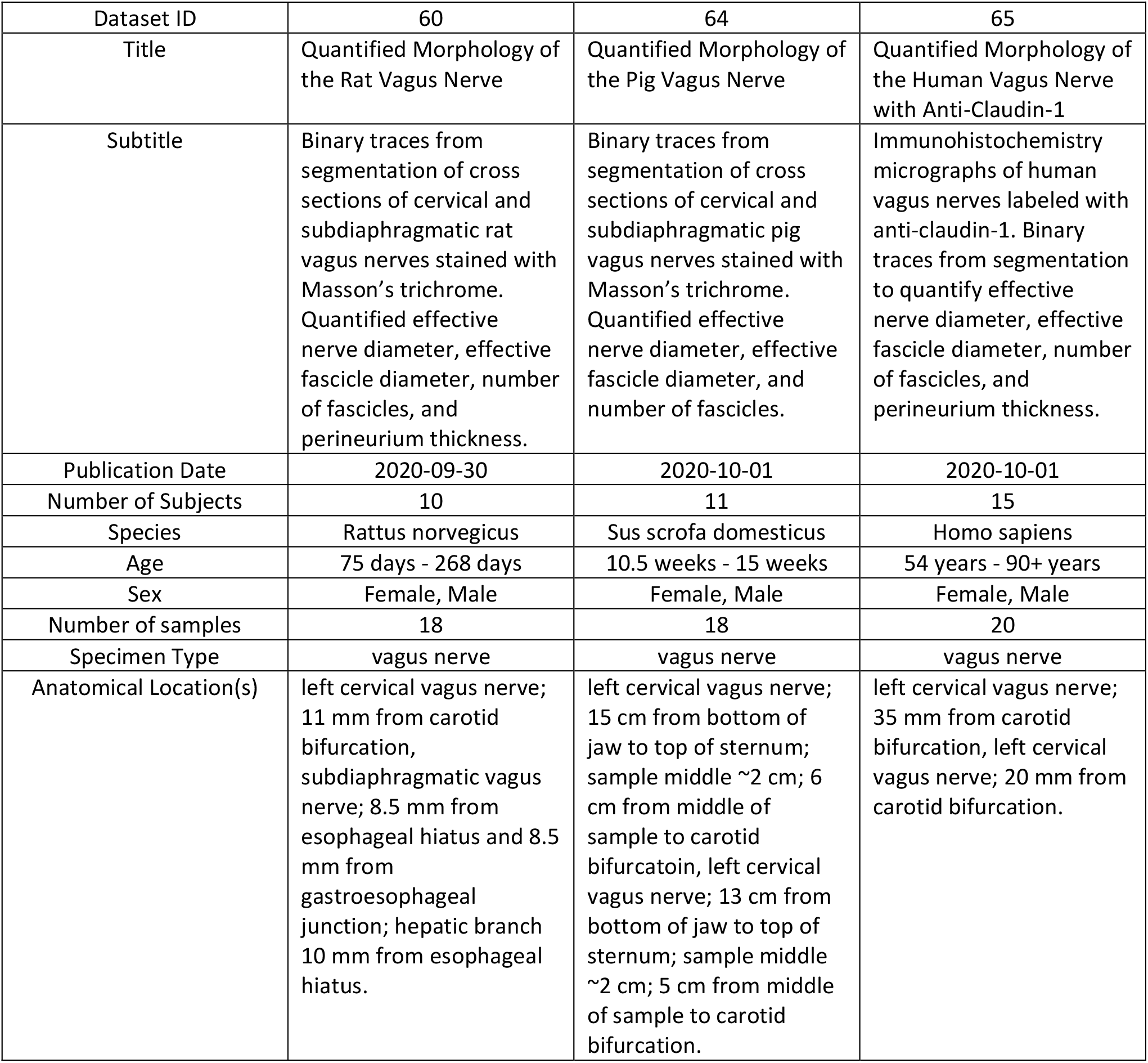
KnowMore Summary Table output for the three datasets in our use case.

| Dataset ID | 60 | 64 | 65 |
| --- | --- | --- | --- |
| Title | Quantified Morphology of the Rat Vagus Nerve | Quantified Morphology of the Pig Vagus Nerve | Quantified Morphology of the Human Vagus Nerve with Anti-Claudin-1 |
| Subtitle | Binary traces from segmentation of cross sections of cervical and subdiaphragmatic rat vagus nerves stained with Masson's trichrome. Quantified effective nerve diameter, effective fascicle diameter, number of fascicles, and perineurium thickness. | Binary traces from segmentation of cross sections of cervical and subdiaphragmatic pig vagus nerves stained with Masson's trichrome. Quantified effective nerve diameter, effective fascicle diameter, and number of fascicles. | Immunohistochemistry micrographs of human vagus nerves labeled with anti-claudin-1. Binary traces from segmentation to quantify effective nerve diameter, effective fascicle diameter, number of fascicles, and perineurium thickness. |
| Publication Date | 2020-09-30 | 2020-10-01 | 2020-10-01 |
| Number of Subjects | 10 | 11 | 15 |
| Species | Rattus norvegicus | Sus scrofa domesticus | Homo sapiens |
| Age | 75 days - 268 days | 10.5 weeks - 15 weeks | 54 years - 90+ years |
| Sex | Female, Male | Female, Male | Female, Male |
| Number of samples | 18 | 18 | 20 |
| Specimen Type | vagus nerve | vagus nerve | vagus nerve |
| Anatomical Location(s) | left cervical vagus nerve; 11 mm from carotid bifurcation, subdiaphragmatic vagus nerve; 8.5 mm from esophageal hiatus and 8.5 mm from gastroesophageal junction; hepatic branch 10 mm from esophageal hiatus. | left cervical vagus nerve; 15 cm from bottom of jaw to top of sternum; sample middle ~2 cm; 6 cm from middle of sample to carotid bifurcation, left cervical vagus nerve; 13 cm from bottom of jaw to top of sternum; sample middle ~2 cm; 5 cm from middle of sample to carotid bifurcation. | left cervical vagus nerve; 35 mm from carotid bifurcation, left cervical vagus nerve; 20 mm from carotid bifurcation. |

#### Common Keywords

The Common Keywords figure provides a graphical depiction of words that show up multiple times across the selected datasets (Figure 4). This size of the word in the image provides a visual representation of the weight (or frequency) of that word across the datasets. Not surprisingly, “nerve” is a large word as it shows up many times. Many other keywords highlight the quantified morphology across the datasets (diameter, cross-sectional area, fascicle, etc.).

**Figure 4.**
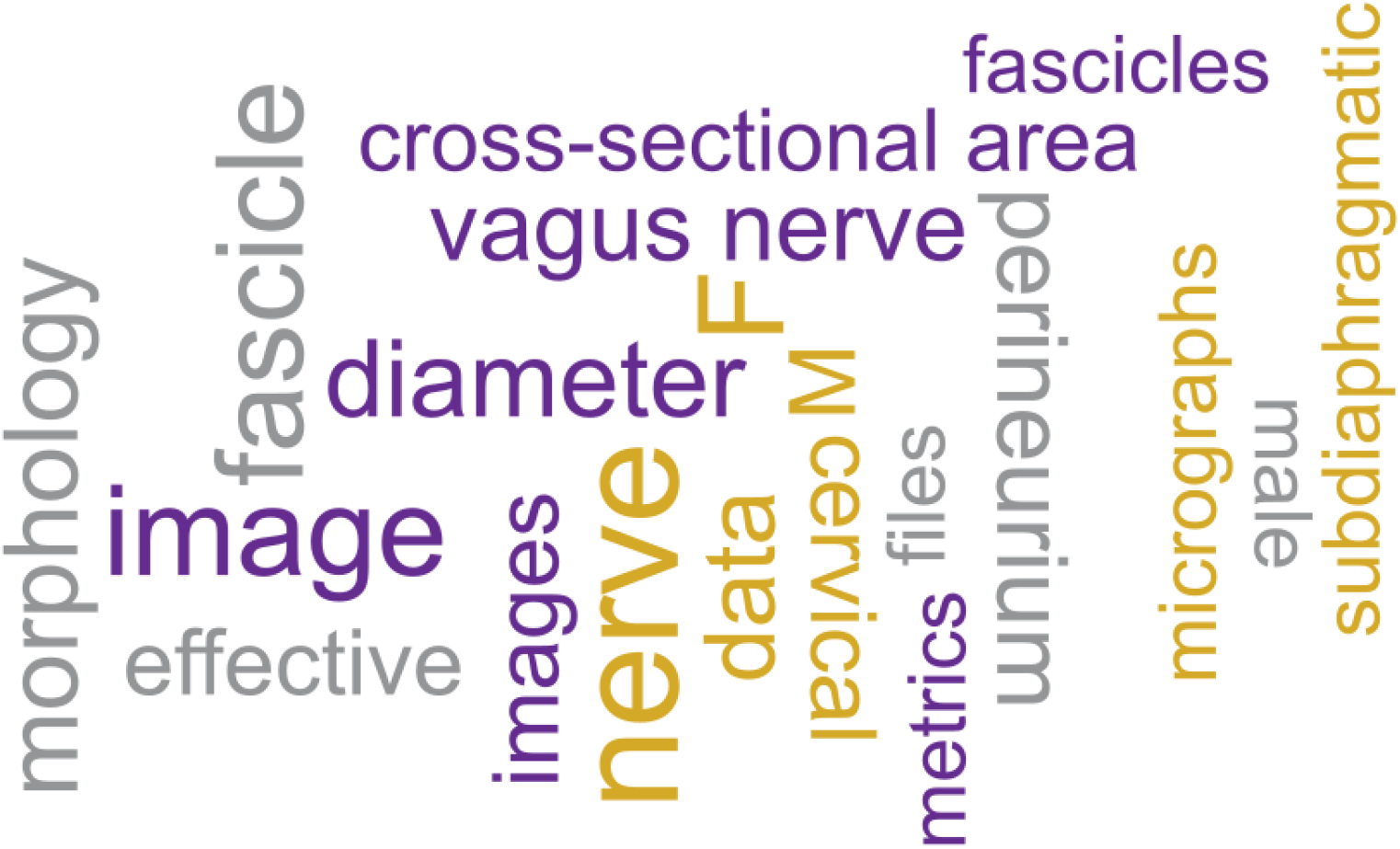
Common Keywords output for the three datasets in our use case.

#### Correlation Matrix and Abstract

KnowMore generates a heatmap illustrating the correlation between the studies based on the words used in the text of these studies (Figure 5). This figure can guide the user in selecting highly correlated studies or eliminating studies that do not correlate well. Additionally, KnowMore generates a combined abstract that provides an overview of all datasets included in the study.

**Figure 5.**
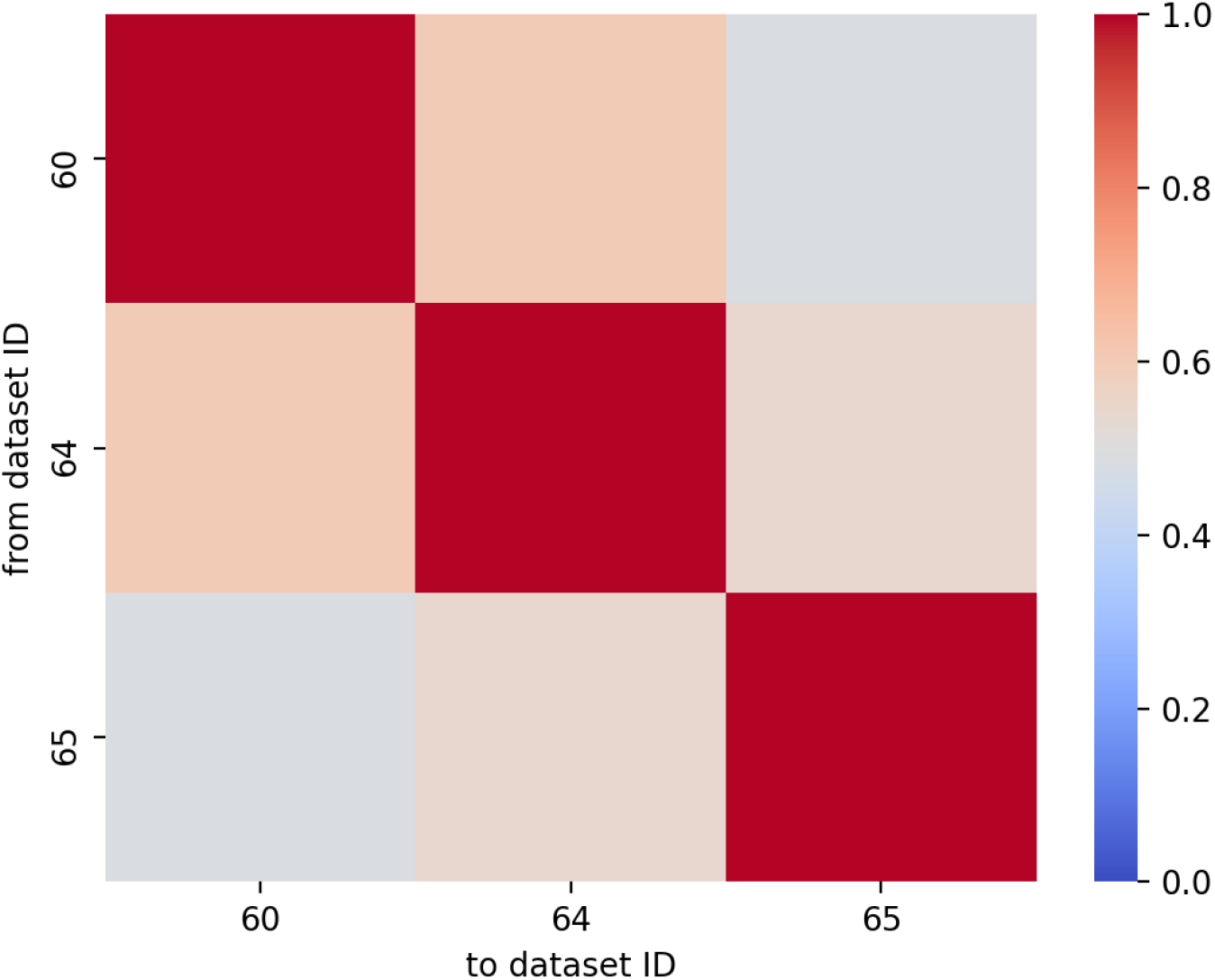
Correlation of the words used to describe the three datasets in our use case.

#### Data Plots

For this use case, KnowMore also generates 20 scatter plots. Due to time constraints, these plots are currently generated in the backend as png files and then displayed in the front-end. Data points are color-coded to each dataset. Each axis is labeled with the variable being plotted. The variable name is obtained directly from the datasets. Three of the plots are presented here (Figure 6). Plot 3.4 reveals that pigs contain more fascicles in their vagus nerves than humans do, and humans contain more fascicles than rats. Plot 3.4 also reveals that pigs and rats have similar variability (spread) in their fascicle diameters whereas humans have a greater spread in their fascicle diameters. Finally, Plot 3.4 illustrates that humans can have larger fascicles than pigs. Plot 3.5 reveals that humans and pigs have similar-sized nerves, though pigs may, on average, have larger nerves. Plot 3.5 also reveals that the number of fascicles in the nerve may tend to be greater for nerves of larger diameter within each species. That is, there appears to be a positive correlation between the number of fascicles in the nerve and the diameter of the nerve. However, Plot 4.5 suggests that there may not be a trend between the fascicle diameter and the nerve diameter. Although these findings have been previously reported in some form^24^, the Data Plots can become a very useful tool in helping researchers quickly understand the underlying data across multiple datasets.

**Figure 6.**
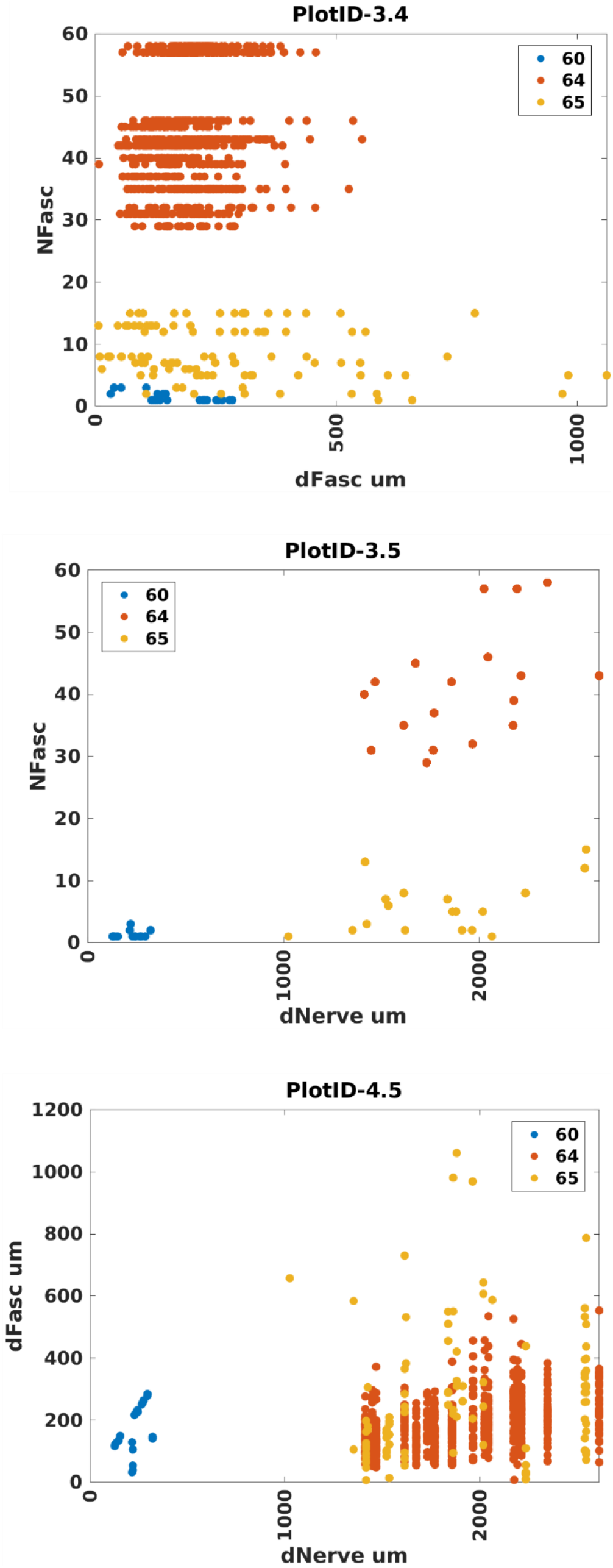
Three selected KnowMore Data Plots created from the three datasets in our use case.

### Conclusions and next steps

#### Potential for this tool

In a few clicks to select datasets, KnowMore can provide both a high-level metanalysis and a granular comparison across two or more datasets on the SPARC portal. KnowMore outputs result at several levels depending on the needs of the researcher. One can quickly determine personnel, institutional, and funding relationships between datasets, and generate an overview of subjects included in the datasets and the techniques used to obtain data. Finally, if data are suitable for plotting, plots can reveal relationships within and across the studies that may reveal larger trends or help the researcher choose or eliminate particular datasets for more detailed analysis.

#### Challenges

SPARC has done an excellent job of standardizing the metadata associated with a study, and, as such, most of the KnowMore output is available across any selected studies. However, SPARC has not enforced standardization for tabular data. As such, the Data Plot output of KnowMore is currently limited to datasets that contain identical variable names and formats. This is an uncommon occurrence across datasets. Data can currently be stored in any number of formats. KnowMore’s Data Plot currently requires data to be stored in a MATLAB .mat file due to our use case, but this could be expanded to several other file formats. It would be preferable from a programming perspective if all data formats and variable naming are consistent, however, within MATLAB alone, data can be stored in multiple formats. Data may be stored in vectors/matrices; cells; cell arrays of vectors, matrices or more cells; structures; or tables, among other formats. Even small differences in variable names such as NerveDiam versus NerveDiameter versus DiameterOfNerve are not immediately reconcilable, though NLP may alleviate such inconsistencies. Without unified variable naming, comparisons across datasets become very challenging. Inconsistent variable names are not the only challenge, however. Even if variable names are identical, the values stored for that variable may be different from study to study. Without unified data types, comparisons across datasets become very challenging. To make the KnowMore Data Plot tool universal we propose standardization of commonly used variable names, data formats, data types, and data units. We also recommend the inclusion of key pieces of information that describe the data in the metadata. We have submitted these recommendations to SPARC and a copy of the document is available in our GitHub repository. This may require a significant amount of effort to convert previously uploaded datasets but should not put an exceptional burden on new studies. Data standardization across the SPARC platform would make the data ready for much broader analysis using more sophisticated big data tools that could provide insights that are otherwise obscured or not readily accessible.

#### Future directions

Currently, the discovery process takes several minutes to run and display the visualization items (about 5 min for the use case). To improve performance, we suggest using multi-threading in the Python script; moving the .mat file processing directly into the Python script; collecting all required raw data (e.g., text) when a dataset is uploaded (e.g. save it in the metadata.json file) and even pre-process it (clean the text) so it is readily available during our discovery process. Image clustering components can be included in the future as well as any other visualization items that are deemed useful to the user. If the above-mentioned challenges with tabular data are addressed, the Data Plots feature of KnowMore can be generalized to work with any datasets.

The SPARC data ecosystem that is built to deliver FAIR datasets, provides a unique opportunity to automate knowledge discovery across datasets. During this project, we leveraged that ecosystem to demonstrated what can be achieved to increase the speed and convenience of discoveries across SPARC datasets. The tool we have developed is a statement of the power of FAIR practices and the effort of SPARC in that regard. We believe that we have only scratched the surface during the Codeathon and the opportunities are yet immense.

## Software availability

Source code available from: https://github.com/SPARC-FAIR-Codeathon/KnowMore^12^

Archived source code at the time of publication: https://doi.org/10.5281/zenodo.5137255^28^

License: MIT

The repository and archive both contain detailed information for using the source code. They also contain a copy of our recommendation to SPARC for standardizing tabular data.

## Acknowledgments

We would like to thank the NIH SPARC Program and the SPARC Data Resource Center (DRC) teams for organizing the 2021 SPARC FAIR Codeathon. We would also like to thank the DRC teams for their guidance and help during this Codeathon.

